# Bidirectional changes in seabird populations drive terrestrial ecosystem function and biodiversity

**DOI:** 10.64898/2026.10.09.757862

**Authors:** Matthew Guy, David Johnson, Tancredi Caruso, Kirsty Park, Lucy Gilbert

## Abstract

Seabird populations fluctuate dramatically in response to perturbations and human activity, such as the current highly pathogenic avian influenza (HPAI) panzootic and fishery practices. Seabirds vector marine nutrients into terrestrial and coastal environments causing cascading effects through ecosystems, but quantifying the inter-related mechanisms underpinning marine-terrestrial cascades in response to changes in seabird populations is challenging. Here we simulate decreases and increases in seabird density, as have occurred due to HPAI and fishing practices respectively, through turf transplant experiments and stratified surveys to quantify marine nutrient cascades and show how they regulate terrestrial ecosystem processes and biodiversity. We use structural equation modelling to quantify the relative strengths of each hypothesised mechanistic pathway in the ecological network. This analysis identified that seabirds shape terrestrial plant and invertebrate biodiversity primarily indirectly, through modifying soil nutrient availability. Our findings highlight the ecosystem changes that are likely occurring, and the pathways through which they operate, due to acute declines and chronic increases in seabirds caused by HPAI, fishing practices or other drivers.

**Significance statement:** We simulate increases and decreases in seabird populations to test their impact on terrestrial ecosystem functioning and biodiversity in Atlantic islands. We show using stratified surveys, modelling and manipulation experiments that invertebrate community diversity and ecosystem processes are driven by cascades of ocean-derived nutrients into soils and through plants via deposition of pellets and faeces from inland-nesting seabirds. Our findings demonstrate how catastrophic decreases in seabird populations, as has occurred due to the avian influenza panzootic, and increases due to marine fisheries, drive terrestrial ecosystem function and biodiversity.

## Introduction

The current highly pathogenic avian influenza (HPAI) panzootic has caused catastrophic declines in many bird populations globally (*1–3*). These declines have potential for wider ecosystem consequences because, as highly mobile species, birds can transport resources such as nutrients between otherwise separate ecosystems (*4*). The flow of resources between ecosystems (*5*) can be a major driver of biodiversity and ecosystem function (*6*). Ecosystems as different as marine and terrestrial can be coupled to form meta-ecosystems by vectors such as seabirds and pinnipeds that transfer nutrients to otherwise nutrient-poor environments (*4,7*), and even back again, to form circular seabird economies (*8*). This process means that perturbations in vector populations can alter the flow of resources to and from marine and terrestrial environments, affecting ecosystem functions, as has been demonstrated by the introduction (*9*) and removal (*10*) of non-native mammals that predate seabirds.

However, ecological networks are complex, comprising interacting direct and indirect pathways between components of a meta-ecosystem, and teasing these apart can be extremely challenging. Major gaps in understanding remain as to the relative strengths of these mechanistic pathways through which meta-ecosystem functioning responds to disturbances. For example, such pathways may operate directly or indirectly, additively or antagonistically and can occur either slowly or abruptly, either promoting or suppressing resource availability (*11,12*).

Mobile vectors linking marine and terrestrial environments may be subject to factors that either increase or decrease their population densities. For example, non-native mammalian predators and marine fishery practices, management and policies can have profound impacts on seabird populations (*9,10,13,14*). While fisheries can have strong negative impacts through several mechanisms such as reducing prey availability or directly killing seabirds in fishing gear (*15*), they can also directly increase food availability by discarding fish at sea (*16*). Fishery discards are important sources of food for many seabird species globally (*17*), including great skuas *Stercorarius skua* in the northeast Atlantic (*18*), and have been a crucial factor contributing to substantial population growth of great skuas and other seabirds through the 1900s (*19*). In stark contrast, the populations of many seabirds globally have suddenly declined significantly due to the global panzootic of HPAI (*1–3*). Great skuas in the northeast Atlantic suffered particularly acute collapse due to mass mortality from HPAI between 2021 and 2023: the largest and most globally significant colonies of great skuas in the archipelagos of Shetland and St Kilda in Scotland suffered reductions of around 80-85% of their populations in a period of less than two years (*20*).

Resolving the extent to which, and the mechanisms for how, bi-directional drivers of seabird populations influence marine-terrestrial interactions is vital to better understand meta-ecosystem functioning and for predicting the wider ecosystem effects of both abrupt and chronic perturbations. These gaps are particularly important for islands and coastal ecosystems because they support disproportionate global biodiversity by land area (*21*) and are more susceptible to be impacted by perturbations (*22*). In particular, many bird species, including great skuas, nest inland on nutrient-limited, carbon rich peatlands that are likely sensitive to sustained long-term additions of nutrients (*23*) but may be equally sensitive to rapid cessation of nutrient inputs.

Filling these gaps in knowledge will allow us to understand and predict the cascading acute impacts of disease panzootics on wider meta-ecosystem functionality and the chronic impacts of changes in the marine ecosystem, including marine fisheries management, on terrestrial communities. By teasing apart the strengths of the direct and indirect mechanistic pathways to these impacts, our approach is directly applicable to other marine-terrestrial meta-ecosystems and to other acute and chronic bidirectional drivers of nutrient transport into soils.

Controlled field experiments that simulate changes in vector populations, and advanced statistical methods that can quantify the influences of the hypothesised pathways in complex networks, would enable us to predict the impact of perturbations in vectors on ecosystem function as well as identify the mechanisms. Here, we simulate acute, dramatic changes in seabird density, as has occurred due to HPAI, by conducting *in situ* controlled manipulation experiments where turfs were reciprocally transplanted between areas of high and low skua density (Supplementary Fig. S3). We make inference about chronic, long-term changes or differences in seabird density through replicated stratified surveys using areas of contrasting great skua density (Fig. 1A, B; Supplementary Fig. 1, 2). We measured a suite of soil, plant and invertebrate parameters and applied structural equation modelling (SEM) to quantify the strengths of each hypothesised mechanistic pathway shaping the impacts of changing seabird density on ecosystem functioning and biodiversity. We quantified the amount of marine nutrients that seabirds transport to a terrestrial ecosystem, and traced these through soil and plants using stable isotope analysis to confirm that soil and plant nutrients are marine-derived and strongly correlated with seabird density. We show that seabird density increases litter decomposition rates, radically changes plant and invertebrate community composition and increases species richness of carabid beetles. The SEM identified that, although direct effects are in evidence, the strongest mechanistic pathways explaining these effects operate indirectly via soil nutrients that alter plant communities and biodiversity, that in turn impact invertebrates. The manipulation experiment demonstrated significant rapid effects of a sudden, dramatic change in seabird density on plant communities and nutrients within 2.5 years of the event.

**Fig. 1.**
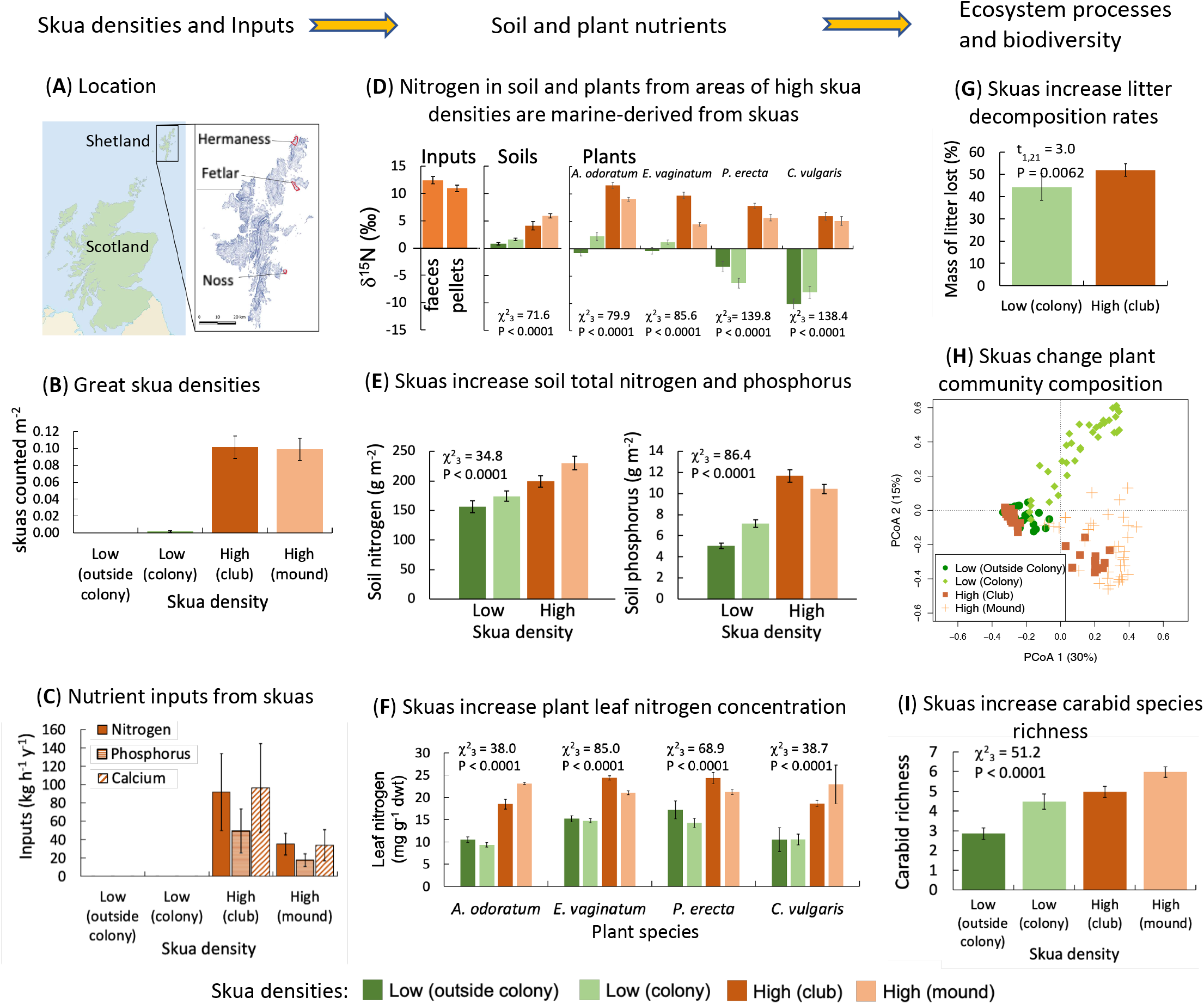
Marine derived nutrients cascade through ecosystems to affect terrestrial biodiversity and function depending on seabird density. (**A**) Location of experimental sites in the Shetland archipelago. (**B**) Great skua densities were significantly greater in high density compared low density sites located both within and outside colonies. Areas with high compared to low great skua density had: (**C**) greater inputs of nutrients from great skua faeces and pellets, (**D**) significantly different d^15^N isotope ratios in plants and soils, which were also closer to those in great skua faeces and pellets, (**E**) significantly greater concentrations of nitrogen and phosphorus in soil, (**F**) significantly greater leaf nitrogen concentrations in all focal plant species, (**G**) significantly faster rates of litter decomposition, (**H**) distinct plant community compositions, and (**I**) more diverse carabid communities (means ± SEM).

## Results and Discussion

### Marine nutrients cascade through terrestrial ecosystems

We identified areas in the Shetland archipelago where great skua densities were categorised as either high or low. High density areas comprised ‘club sites’, which are areas within colonies where adult and immature non-breeding birds congregate, and ‘mounds’ where breeding individuals stand to overlook their breeding territory. Areas categorised as low density comprised most of the ground within great skua colonies that is not within core territories near nests. Additional low density areas were located outside colonies that had similar geology, habitat and topography to the closest colony. Observational data confirmed that great skuas were absent outside colonies and infrequent within the colony low density areas compared to club sites and mounds, which supported similar high densities of birds (Fig. 1B). We quantified inputs of nitrogen (N), phosphorus (P) and calcium (Ca) from faeces and regurgitated pellets of great skuas across all areas (Fig. S2) which confirmed that low density areas received no measurable inputs of N, P or Ca, whereas high density areas received substantial inputs (Fig. 1C; *P* <0.001 for N, P and Ca). For example, club sites received around 90 kg N ha^-1^ y^-1^ and 45 kg P ha^-1^ y^-1^ (Fig. 1C). These inputs of N are double the highest modelled rates of wet and dry atmospheric N deposition in the UK and far exceed the average background rates of atmospheric N deposition in Shetland (10 kg N ha^-1^ y^-1^; *24*). Moreover, the substantial inputs of P alongside N are likely to alleviate co-limitation on plant-soil responses and thus generate greater impacts on ecosystem functioning than from inputs of N alone (*25*).

To test whether it is the marine-derived N from great skua inputs, rather than some other source of N, that cascades through the terrestrial ecosystem, we analysed stable isotope ratios of N (δ^15^N) in great skua faeces and regurgitated pellets, soils and the shoots of plants from four plant functional groups (Fig. 1D). Soils from high density great skua areas had significantly more positive δ^15^N, closer to the δ^15^N in faeces and pellets, compared to soils from low density areas. Similarly, plants growing in high great skua density areas had substantially higher shoot δ^15^N than plants growing in low density areas (10 - 16 ‰ shift; Fig. 1D). The significant positive shifts in δ^15^N in shoots in response to great skua density were consistent across plant species regardless of the functional type they represented (grasses, sedges, forbs, shrubs). The greatest shift (∼16‰) occurred in the slow-growing shrub heather, *Calluna vulgaris* (Fig. 1D). The magnitude of this shift in δ^15^N is extreme, and even greater than changes seen in response to long-term exposure to 64 kg N ha^-1^ y^-1^ of simulated wet deposition (*26*) and large gradients in N deposition in urban landscapes (*27*). The shifts in δ^15^N in response to high skua density were mirrored by increases in total N and P concentrations in soil (Fig. 1E) and N in plant leaves (Fig. 1F). The nutrient and stable isotope data demonstrate that substantial quantities of growth-limiting nutrients from the marine environment vectored by great skuas cascade through terrestrial soils and vegetation and generate biologically significant changes in the chemistry of peatland soils and plants. While nutrient transfer from seabirds to plants and soils is well documented (*4*), previous studies are typically from colonies containing seabird densities orders of magnitude greater than those in the great skua colonies studied here where nests are tens of metres apart and bird densities are generally low.

### Marine nutrients drive biodiversity and ecosystem function

We assessed the consequences of these nutrient cascades on plant litter decomposition, a key ecosystem process, and plant and invertebrate diversity, which regulate numerous critical functions, such primary productivity and stability (*28,29*).

Litter from high great skua density club sites had N concentrations that were 65 % greater, and P concentrations that were 250 % greater, than litter from low great skua density sites outside the colonies (Table S4). The carbon (C) content of litter did not differ between high and low density sites, so club sites had significantly lower C:N and C:P ratios (Table S4), which are crucial metrics controlling decomposition (*30*).

Furthermore, litter from club sites was significantly enriched in δ^15^N (Table S4) indicating transfer of N by great skuas from marine derived resources. Because of these changes in chemistry, litter from high great skua density areas decomposed significantly (17%) faster than litter from low density areas (Fig. 1G). These findings suggest litter decomposition is likely a key pathway of marine nutrient inputs into soil food webs that promotes recycling of nutrients within the terrestrial environment.

Plant community composition was significantly different between low and high great skua density areas, with the first two axes of a principal coordinates analysis accounting for 45% of variation in the data (Fig. 1H). High great skua density areas tended to be characterised by greater abundance of species of plants exhibiting ruderal life strategies (including *Sagina procumbens*, *Juncus bufonius* and *Stellaria media*) and, at one location (Noss), a negative association with stress tolerant life strategies (including *Carex nigra*, *Anthoxanthum odoratum*, *Luzula multiflora*, *Potentilla erecta* and *Galium saxatile*; Fig. S4, Table S1). Ruderal life strategies are associated with fast growth rates and resource allocation to seed production, allowing them to outcompete other species in low nutrient, disturbed environments (*31*), including within gull colonies in the Mediterranean and in North America (*32–34*). Although we were unable to disentangle effects of physical disturbance from nutrient addition, fertilisation from seabirds alone has been shown to be sufficient to cause large-scale change in vegetation (*10*). In addition, great skuas form low density colonies and, unlike burrowing seabirds, any physical disturbance would be restricted to modest damage, such as trampling, to the above-ground parts of plants. Thus, we expect that the dominant mechanism underpinning our observations is from nutrient transfer in faeces and pellets.

We also quantified the effects of great skua density on terrestrial invertebrate communities by identifying a total of 47,878 individual invertebrate specimens to order level from pitfall traps across all sites. Multivariate analysis showed that the invertebrate order community was significantly influenced by great skua density (Fig. S5, Table S2; score_[3,144]_ = 269.7, *P* <0.002). Of the 18 orders identified, high great skua density sites were characterised particularly by greater abundances of Diptera (flies), Collembola (springtails), Hemiptera (true bugs) and Coleoptera (beetles) than were low density sites (Fig. S5, Table S2).

A total of 5324 (11%) of the invertebrate specimens were identified as beetles comprising 14 families, and the abundance of individuals in this order was positively correlated with those from 10 other orders (Fig. S6), suggesting that beetle abundance is a useful indicator of invertebrate abundance in this ecosystem. Carabids (ground beetles) were the most abundant beetle family identified from pitfall traps, accounting for just under half of all Coleoptera (2465 specimens). Carabids across all great skua colonies comprised 24 species, and the abundances of 54% (13 out of 24) of the carabid species were significantly influenced by great skua density (Table S3), resulting in greater carabid species richness in high skua density areas (Fig. 1I). High skua density sites supported particularly greater abundances of *Loricera pilicornis*, *Agonum fuliginosum, Pterostichus rhaeticus*/*nigrita*, *P. strenuus* and *P. diligens* than did low skua density sites, whereas low skua density colony sites had more *P. adstrictus* (Fig. S7).

Carabids are a diverse and widespread family, comprising mainly top-invertebrate predators, that are frequently used as bioindicators of ecosystem health because of their ecological importance and sensitivity to disturbance (*34*). Our data demonstrate their responsiveness to marine-derived nutrients. More generally, invertebrates drive numerous critical ecosystem functions, such as pest control, pollination and relationships between plant species richness and ecosystem processes (*36,37*), but there is less knowledge of the mechanisms driving these effects (*38*). Our findings demonstrate how marine nutrient transfer from seabirds is a key driver of diversity of above-ground terrestrial invertebrates that span trophic levels (i.e. vertical diversity; *39*).

### Pathways through which great skuas influence biodiversity

We hypothesised the mechanistic pathways through which we expected great skua density to drive plant and carabid species richness, and used structural equation modelling (SEM) to quantify the statistical support for each component of these pathways (Fig. 2). Several pathways were identified through which great skua density influenced plant species richness, including negative and opposing positive effects. The strongest pathway between great skua density and plant species richness operated positively through a strong increase in soil nutrient concentrations which shifted plant community composition, while there was an additional weak direct positive effect of great skua density on plant community composition, perhaps caused by physical damage. The shift in plant community composition was characterised by an increase in ruderal species, particularly *Holcus lanatus* (the grass Yorkshire fog) and *Deschampsia flexuosa* (wavy hair-grass) and a decline in *C. vulgaris* (heather), the overall manifestation being a strong increase in plant species richness (Fig. 2). In addition, there was a direct negative effect of great skua density on plant species richness, which may be related to physical damage to plants or foliar uptake from direct deposition of nutrient-rich faeces onto leaves; and a slightly stronger negative effect operating indirectly via the large increase in soil nutrients which then reduced plant species richness (Fig. 2). Great skuas shaped carabid species richness via pathways which acted through plant community composition and additionally via plant species richness (Fig. 2). These measurements from areas of contrasting great skua density combined with the SEM analyses demonstrate how transfer of marine nutrients onto land drive key ecosystem processes and connectivity among terrestrial biodiversity groups across multiple trophic levels.

**Fig. 2.**
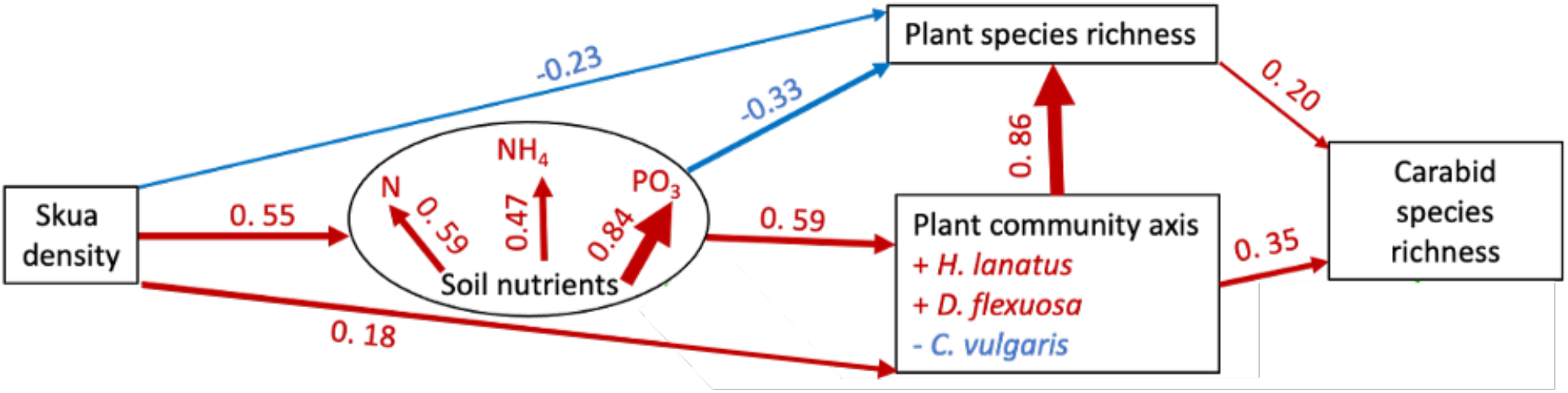
Structural equation model of the direct and indirect effects of great skua density on plant and carabid communities. Red = positive associations; blue = negative associations. The values above the lines are path coefficients and the width of the arrow relates to the size of the coefficient. The indirect effects were hypothesized to be exerted through the impact of great skuas on soil nutrients, which were modelled as a latent variable (circle) in which the best fitting soil variables (extractable ammonium, nitrate and phosphate concentrations) were used. The modelling selection process supported the inclusion of both plant species richness and composition (represented by the first PCoA axis of the ordination reported in Fig. 1H) and in using carabid species richness as a description of the insect assemblage. Direct effects of great skua on carabid communities were not supported while indirect effects via the changes that great skua caused on the plant community were supported. Overall, the SEM supports the hypothesis that variation in great skua density cascades through the plant community both directly and indirectly (with the two effects having a similar magnitude) and indirectly on the rest of the biota (in the model represented by the top predator group of carabid beetles). Global fit metrics: χ^2^ = 9.27 (d.f. = 9; *P* = 0.41); comparative fit index = 0.999; root-mean square of approximation = 0.015.

These findings have clear implications for how changes in the marine ecosystem, fisheries management, disease or other factors that influence seabird populations and marine-terrestrial nutrient flow, could affect terrestrial ecosystems. The pathway coefficients in our SEM linking great skua density to soil nutrients and other trophic levels, in addition to the findings from our surveys across areas of contrasting great skua density, strongly suggest that changes in great skua density will cause trophic cascades through the terrestrial ecosystem. Experiments to simulate such changes in great skua density would be a robust way of empirically testing these cascading effects.

### Empirical experiments to simulate impacts of changes in great skua densities

We therefore undertook an *in situ* manipulation experiment (Fig. S3) to simulate two important types of bi-directional changes in great skua density that have occurred in nature to gain greater insight into causation. The first change concerns chronic increases in great skua densities. These increases have averaged 525% over all UK colonies since the 1980s (*19*) and are linked to fisheries practices, especially discarding fish at sea that provided a large proportion of the great skua diet during the breeding season (*18*). The second change concerns the acute, catastrophic (80-85%) collapse in great skua densities that occurred 2021-2023 due to mass mortality from the global epizootic outbreak of high pathogenic avian influenza (*2,20*). To test the hypothesis that bi-directional changes in great skua density will impact the terrestrial ecosystem, intact turfs were transplanted from areas of high great skua density to areas of low great skua density and *vice versa* and, to account for possible effects of disturbance from the transplant process, control turfs were also removed but returned to their original environment. The turfs were encased in free-draining boxes that prevented lateral movement of nutrients to and from the surrounding landscape. We monitored vegetation communities over three growing seasons and analysed plant tissue N and P concentrations in four plant species each from a different functional group (a sedge *Eriophorum vaginatum*, a grass *A. odoratum*, a forb *P. erecta*, and a shrub *C. vulgaris*).

#### Simulated increases in great skua density influence terrestrial ecosystems

We found that turfs transplanted to areas of high great skua density led to a rapid change in plant community composition. Specifically, plant communities in turfs that were transplanted from low to high great skua density areas became more similar to vegetation growing in high great skua densities (Fig. 3A). In particular, increased great skua density led to a rapid decline in dwarf shrub species and an increase in graminoid (grasses) cover, especially by *A. odoratum*. Moreover, the concentration of N in leaf tissues increased significantly in two of the four (*E. vaginatum* and *P. erecta*) focal plant species in turfs transplanted from low to high great skua density environments (Fig. 3B), with similar trends being observed in the other species, and for leaf P concentrations (Table S5). Our combined results suggest that the increase in great skua abundance since the 1980s would have strengthened the coupling between marine and terrestrial ecosystems, greatly increasing marine nutrient transfer into soils, leading to rapid changes in terrestrial nutrient cycling, litter decomposition rates and plant and invertebrate communities as the colonies grew and more non-breeding birds attended club sites.

**Fig. 3.**
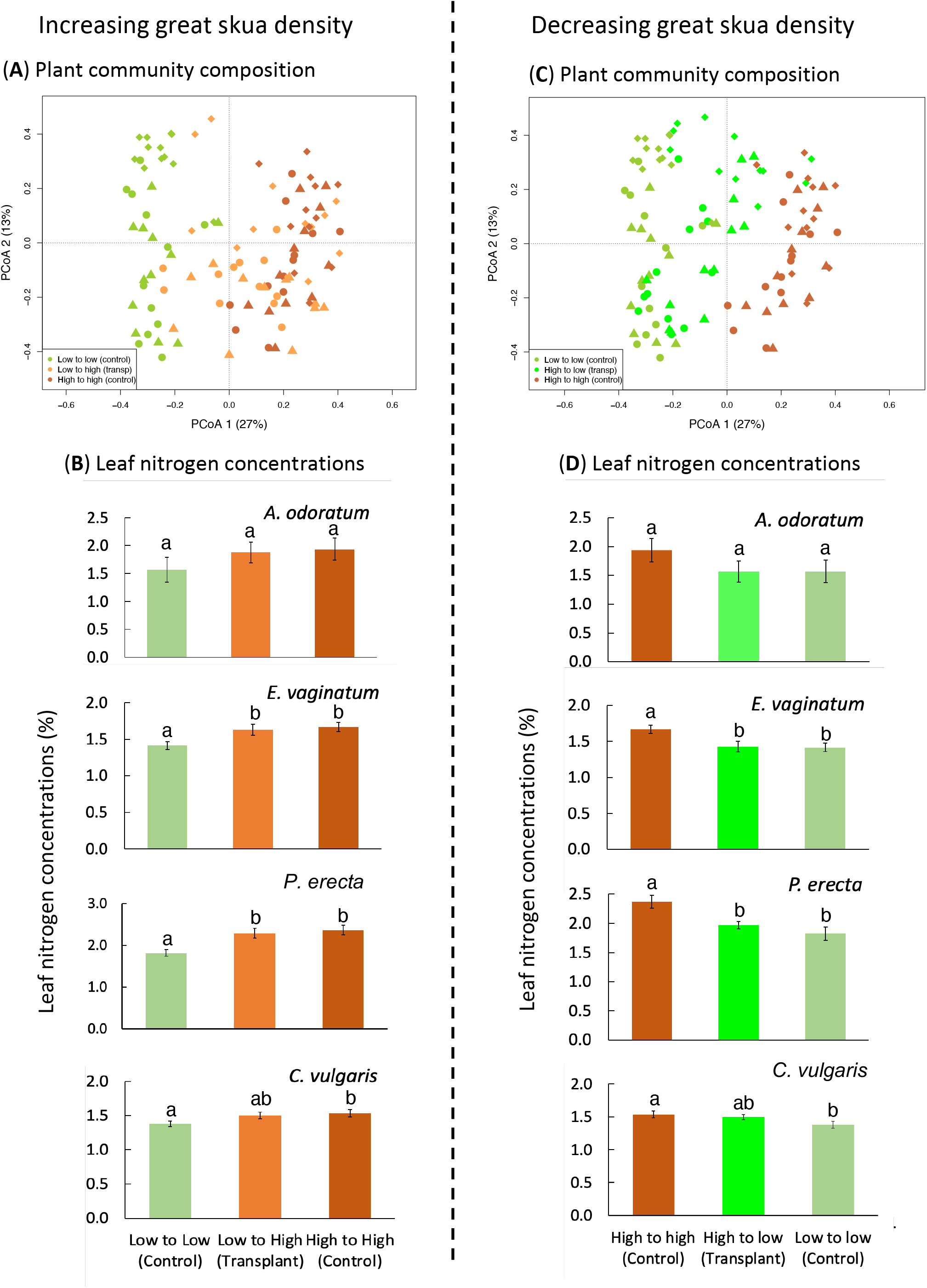
Simulated increases and decreases in great skua density reconfigured plant communities and leaf nitrogen concentrations. Simulated increases in great skua density through transplantation of turfs (**A**) shifted plant communities and (**B**) leaf nitrogen concentrations towards those found in high great skua density areas. Simulated decreases in great skua density through transplantation of turfs (**C**) shifted plant communities and (**D**) leaf nitrogen concentrations towards those found in low great skua density areas (means ± SEM). Bars sharing a letter are not significantly different (*P* >0.05). Turfs were either extracted and replaced in their original skua density (high great skua density controls: brown bars and symbols; low great skua density control turfs: sage green bars and symbols) or transplanted to a new great skua density (low to high density transplants: orange symbols and bars; high to low transplants: bright green symbols and bars).

#### Simulated collapses in great skua density influences terrestrial ecosystems

When turfs were moved from high to low great skua density areas, there was also a substantial change in plant community composition after only 2 years, with plant communities becoming similar to those growing in low great skua density areas (Fig. 3C). This finding demonstrates that great skuas couple the marine and terrestrial ecosystem both spatially and temporally, with surprisingly rapid effects. Furthermore, N concentrations declined significantly in the shoots of *E. vaginatum* and *P. erecta* (Fig. 3D), and similar declines were seen in leaf P concentrations (Table S5), suggesting residual soil N and P pools became depleted over the duration of the 2-year transplant experiment indicating great skuas generate little legacy effects. Other work shows that after local seabird colony declines due to invasion of rats to warm-temperate forested islands, P and to a lesser extent N, can remain at high concentrations in soil even after 50-150 years (*40*). Our results differ and indicate plants utilized residual nutrients that moderated shifts in community composition, and thus our cool-temperate peatland system is responsive over relatively short-timescales (∼2 years) to both increases and decreases in marine-derived nutrient inputs.

The near-cessation of marine nutrient inputs from the massive and rapid loss of the great skua population due to highly pathogenic avian influenza in Shetland between 2021 and 2023 is likely to be sustained for many years. This is because great skuas are long-lived, slow-reproducing seabirds that suffered very high adult mortality so that population recovery is likely to be very slow. We therefore predict a shift in plant communities in the great skua colonies away from ruderal graminoids and towards ericaceous shrubs, that will reduce plant community heterogeneity. We expect that the now very low great skua densities will generate more homogeneous landscapes, likely associated with reduced ecosystem multifunctionality of these nutrient-poor peatlands (*41*). These findings also highlight the potential for disruption of marine-terrestrial linkages globally, not only because highly pathogenic avian influenza continues to cause mass mortalities of seabirds and pinnipeds worldwide, including sub-Antarctic islands, but also because of the widespread links between fisheries practices and seabird foraging (*16*) causing population declines (*42*).

## Conclusions

Here we provide evidence from empirical field-based experiments, supported by surveys and modelling, that the diversity and functioning of island terrestrial environments respond profoundly and rapidly to both increases and collapses in seabird densities. Nearly a third of seabirds worldwide are in danger of extinction (*8*), and these concerns are reflected in the drastic population changes that have occurred in our system and worldwide due to marine fisheries management and the outbreak of a global infectious disease pandemic. These drivers have both chronic and acute catastrophic effects on marine nutrient vectors such as seabirds and pinnipeds. While islands occupy only 6.7% of the land surface, they contain around 20% of Earth’s biodiversity (*21*) and are disproportionally impacted by climate change and other perturbations (*22*). Our findings emphasise the vulnerability of the terrestrial ecosystems of Atlantic islands and other areas globally due to anthropogenic and natural perturbations of vectors of marine resources to land.

## Methods

### Study system and general experimental approaches

Data were collected over three great skua breeding seasons at three large great skua (*Stercorarius skua*) colonies in Shetland, UK, comprising a mix of blanket bog and modified bog: Lamb Hoga peninsula, Fetlar (Fetlar: centred at 60° 34’ 39.77”N, 0° 53’ 33.63”W), Hermaness National Nature Reserve, Unst (Hermaness: centred at 60° 49’ 0.15”N, 0° 53’ 32.40”W) and Isle of Noss National Nature Reserve (Noss: centred at 60° 8’ 22.94”N, 1° 1’ 2.40”W; Supplementary Fig. 1). The vegetation communities at Fetlar and Hermaness have affinities to National Vegetation Classification (NVC) community M19 *Calluna vulgaris – Eriophorum vaginatum* blanket mire blanket mire vegetation, whereas that at Noss has a greater affinity to NVC community M17 *Trichophorum cespitosus – Eriophorum vaginatum* blanket mire and M15 *Trichophorum cespitosus –Erica tetralix* wet heath, as there is a reduced abundance of *Calluna vulgaris* largely due to historic land use (*43*). Both Noss and Hermaness are extensively grazed by sheep.

Great skua abundance across each colony is not homogeneous, but comprises a loose collection of breeding territories, each one containing the nest and at least one raised mound on which the birds stand to observe their territory, which we refer to as “high density – mounds”. In addition, there are one or more discrete areas called club sites where adult and immature non-breeding birds form dense aggregations of between 10 and 200 individuals (*44*); we refer to these as “high density – club”. Most of the land area of a colony away from nests, mounds and club sites is used infrequently by great skuas and is referred to here as the “low density -colony”. In addition, we identified and sampled from areas that were spatially distinct from the colonies, supporting no great skua breeding colony, but which shared similar land-use, soils, geology and distance and orientation to the sea. These areas are referred to as “low density – outside colony”. A total of 12 sampling sites were identified for each treatment type and locations are presented in Supplementary Fig. 1. Mound sampling sites were selected at random from all breeding territories identified within approximately a 1km radius of the club sites (*45*), and club sampling locations were established in a grid formation separated by at least 5m. Low density -colony sampling sites were positioned at approximately 30 m from each mound sampling site on a random bearing, ensuring there were no other nearby mound sites. Low density – outside colony sampling sites were established in a grid formation separated by at least 15m.

We undertook two broad experimental approaches, both of which were reviewed by the Animal Welfare & Ethical Review Body at University of Stirling. They comprised stratified surveys that were extensive sampled across great skua colonies and areas where great skuas were absent to quantify i) nutrient inputs from regurgitated pellets and faeces, ii) soil nutrient status, iii) plant nutrient status, iv) natural abundance stable isotope ratios of faeces and pellets, soils, plant leaves and litter, v) plant community composition, vi) above-ground invertebrate community composition, and vii) rates of litter decomposition. The stratified surveys used high density club and mound sites and low density colony and outside colony (control) sites previously identified.

A second complementary approach used a manipulation study that comprised a reciprocal transplant experiment of intact turfs from areas with high and low great skua density, to simulate the effects of either increasing or decreasing great skua populations on plant community composition and nutrient concentrations. The experiment was established at a high density club site and an adjacent low density -colony area at Hermaness, in a separate area of the colony used for the stratified surveys. In March 2016, pairs of turfs in a block design (n =12; block locations are shown in Supplementary Fig. 3) were cut from club and intra-colony areas and placed into freely draining plastic boxes (35cm x 53cm x 15cm deep). One of the pair was reciprocally transplanted with the corresponding block in the club or intra-colony area, while the other was placed back into the hole from where it was cut, establishing four different treatments: 1) Club to club area (high to high skua density; control); 2) Club to low density colony area (high to low skua density treatment); 3) Low density colony to club area (low to high skua density treatment); 4) Low density colony to low density colony area (low to low skua density; control).

### Quantifying nutrient deposition by great skuas

Recent analysis of pellets has shown that great skuas in our focal colonies have a diet that is almost entirely marine-derived (*46*). To calculate the amount of marine derived nutrients introduced into the terrestrial ecosystem by great skuas, rates of faecal and pellet deposition were measured at high density club and mound sites and low density colony sites at all three colonies (Supplementary Fig. 2). In addition, faecal and pellet samples were collected to measure the nutrient content. Five club sites were selected across the three colonies: two at Hermaness (Fild and North), two at Noss (East and West) and one on Fetlar. Within each club site, sampling locations were established in a grid formation, each one separated by at least 5m. Within colonies, mound sampling sites (n =30) were selected at random from all breeding territories identified within approximately a 1km radius of the club sites. The exception to this was at Hermaness, where several mound sites were approximately 2km south of the nearest club site. Low density colony sample locations were established 20-30m from each mound site on a random bearing (n = 30).

Faecal deposition was quantified in 1m^2^ quadrats and pellets quantified in 4m^2^ quadrats in June 2016. The quadrats were cleared completely of faeces and pellets and after 15 days, faeces and pellets within the quadrats were counted. Where faeces overlapped, a count was approximated based on size, shape and orientation of the outline.

The chemical composition of faeces and pellets were determined in 12 samples of each collected at random from within the quadrats. All additional material, such as grass and soil, were removed from the samples by hand, before each sample was air dried, weighed, and ball-milled. Nitrogen concentrations were quantified using an automated elemental analyser (FlashSmart, Thermo Scientific). Phosphorus and calcium concentrations were determined using a nitric acid microwave digest (MARS 6, CEM) and subsequent analysis using ICP-OES (iCAP 6000, Thermo Scientific).

### Calculating nutrient deposition rates

Nutrient deposition is expressed as mean kg of nutrient per hectare per year. Concentrations of nutrients were converted into grams per dry weight, averaged across the samples and multiplied by respective deposition and scaled-up rates to calculate kg of nutrient deposited per hectare per day. The length of time spent at the breeding colony in a season by breeding pairs and non-breeding individuals was calculated based on dates of arrival and departure (*44*).

In addition, the total amount of marine nitrogen and phosphorus transferred in a breeding season by great skuas was estimated. The areas of all club sites were measured using geographical information system (GIS) and the area of a sub-set of 12 mounds were measured at each colony. To estimate the total area of mounds at each colony, mean mound areas were multiplied by the number of breeding pairs recorded during the Seabird 2000 census of the UK (*47*). This figure was then used to convert deposition rates into total deposition.

### Plant and soil sampling

In July 2017 and August 2018, plant samples were collected from half of the locations used in the stratified surveys (n = 6; sample locations selected randomly), and from all turfs in the manipulation study where the species occurred. Plant samples comprised fresh, above ground vegetative tissue (i.e. not flowering stem) of four functional groups: a sedge: *Eriophorum angustifolium,* a grass: *Anthoxanthum odoratum*, a forb: *Potentilla erecta*, and a shrub: *Calluna vulgaris*. These species were selected because they are ubiquitous across the study area and have contrasting life histories. The plant samples were air dried and ball-milled before chemical analysis.

In June 2017 a single soil core (diameter = 4cm, depth = 10cm) was taken at each sample location in the stratified surveys. The soil cores were returned to the laboratory, sieved to 2mm, air dried, and ground using a pestle and mortar prior to chemical analysis. At each sample location in the stratified surveys, a second core was taken using a bulk density sampler (diameter = 5.4cm, depth = 5.9cm). Soil was air-dried and weighed to determine dry weight per volume of soil. This allows for nutrient concentration in the soil to be converted from mg of nutrient per g of soil to mg of nutrient per m^2^ of soil. This conversion corrects for any influence the physical soil properties may have on resource availability allowing for accurate comparison of samples over a large geographical area.

### Nutrient and stable isotope analysis

The concentration of total nitrogen and phosphorus in soil and plant samples were measured as described for faeces and pellets. Extractable nutrients were also measured in the soil samples. Ammonium and nitrate were extracted by shaking soil in 1M KCl for 1 hour, while phosphate was extracted by shaking soil in 2.5% acetic acid for 2 hours. Nitrogen extracts were stored at −20°C, while phosphate extracts were stored at 4°C, prior to analysis a SEAL AutoAnalyzer 3 constant flow analyser system (Seal Analytical, UK).

Stable isotope analysis was used to determine if the nitrogen in the soil and plant tissue was marine derived. A sub-section of the faecal and pellet samples, plant leaves and soil samples (n = 6) were analysed for the ratio of ^14^N and ^15^N using continuous-flow isotope ratio mass spectrometry (CF-IRMS, Costech ECS 4010 elemental analyser and Thermo Finnigan Delta Plus XP mass spectrometer). Results are expressed in δ^15^N (‰) deviations from the international standards.

### Litter bag study design

Senescing graminoid plant material was collected from a club site and an adjacent area of low density colony at the North Hermaness colony. Litter was air dried, cut to approximately 2cm strips and mixed to produce two bulk samples of litter from high and low density areas. A total of 24 nylon mesh litter bags, measuring 4.5cm x 4.5cm, with a 1mm aperture, were filled with 0.5g of either club (12 litter bags) or control litter (12 litter bags). Litter bags were buried vertically in the top 5cm of soil in a block formation with club litter bags buried at high density club sites and low density colony litter buried in low-density colony areas (n =12) and incubated 24 months. Once harvested, litter was air dried and roots and soil were removed by hand. The litter was weighed to determine mass loss prior to undertaking chemical analysis, as described previously.

### Plant community analyses

The pin drop method was used to quantify the plant species abundance and community composition in both the stratified surveys and manipulation experiment. In the stratified surveys, 60cm^2^ quadrats were randomly placed within sampling locations during July 2017. A total of 16 pins were positioned evenly across the quadrat (15cm spacing). Each vascular plant touching the pin was identified to species level (where possible) and the number of times each species touched the pin recorded. Bryophytes, liverworts and lichens were also recorded, but not identified so are not included in the analysis. The pin touches across the quadrat were summed and used as an objective measure of relative abundance.

The same method was used for the turf transplants; however, 16 pins were placed evenly in a 20cm quadrat (5cm spacing). Quadrats were positioned in the centre of the turfs and away from any other sampling events occurring within the turfs (i.e. soil cores). Vegetation was sampled in early August on three occasions: in 2016, 2017 and 2019 (i.e. 5 months, 17 months and 29 months after transplantation).

### Invertebrate sampling

Invertebrate sampling was carried out using pit-fall trapping. Traps comprised plastic cups (127mm depth x 96mm diameter) buried flush with the soil surface and partially filled a 50% propylene glycol antifreeze solution (diluted with water). In a preliminary study, flooding of traps and interference by great skuas was an issue. To overcome this, two traps were set at least 1m apart within each sampling location. One trap from each site was selected randomly (unless it was the only one still intact) for sample processing. Traps were protected by a clear lid (140mm diameter) held 15mm from the ground by two U shaped pegs. Traps were established in June 2017 and were left open for a period of 4 weeks and emptied at two-week intervals. The trap contents were stored in 70% ethanol, before all individuals were identified to order, all Coleoptera were identified to family and all carabids to species. The exception to this was *Pterosichus rhaeticus/nigrita*, which can be reliably separated only using genetic analysis (*48*). Several summary descriptors and indices were calculated to provide an overview of the carabid community and the effect of great skua use. These comprised the proportion of the total Coleoptera individuals that were carabids, the total number of individual carabids recorded (abundance) and carabid species richness.

### Statistical analysis

#### Stratified surveys

The effect of great skua use on nutrient inputs, isotope ratios, nitrogen and phosphorus concentrations in the soil and plant material, and carabid richness was assessed using mixed effects models using lme4. Great skua density was included as a fixed effect and colony (Fetlar, Hermaness, Noss) as a random effect. Each plant species was assessed separately. The effects of great skua density were assessed using likelihood ratio test and then estimated marginal means were calculated to identify differences between groups. Simple linear model used to assess mass of litter lost predicted by great skua density. A Welch Two Sample t-test was used to assess differences in the mass lost.

Plant community composition was analysed by principal coordinates analysis (PCoA). We use Principal Coordinate Analysis (PCoA, a generalization of Principal Component Analysis or PCA) to describe gradients in plant community composition. Classic PCA implies an underlying Euclidean space, which is generally not appropriate to model distances between communities due to changes in species composition and relative abundance. We, therefore, followed (*49*,*50*) and Hellinger-transformed species density data to work with ecological meaningful distance metrics while retaining the power of ordinations based on the eigendecomposition of the correlation or variance-covariance matrix. We used the R “vegan” packages (*51*) to apply the transformation with the function “decostand” and calculate an eigenvector decomposition on the transformed data, which equates to a PCoA The first axis accounted for a relatively large fraction of total variance due to a large number of strong correlations between species, so the first axis was used in subsequent analyses as a descriptor of major changes in plant community structure. The loadings of the first eigenvector were used to interpret the axis in terms of which species increased and decreased across the sites, and so to interpret the changes in plant species composition associated with the variables under investigation, notably great skua density and soil nutrient concentrations.

The response of individual plant species to the great skua treatments was tested using generalized linear models (GLM) using the MVAbund package, in R (*52*). In the MVabund package, separate GLMs were fitted to each response variable in the analysis using a common set of explanatory variables. Here, the response variables comprised the number of pin touches for each species at each sampling location (i.e. a site by order/species matrix). Great skua treatment, colony and an interaction between the two were included as fixed effects. Score tests (a resampling-based hypothesis test) using PIT-trap resampling were then used to initially make community-level (i.e. all taxa abundances at a sample location combined) and, subsequently, taxon-specific inferences about how the explanatory variables influenced the plant species abundances. This test and the resampling technique was considered most appropriate for these data structures, whilst still taking into account correlations between taxa abundance (*52*).

To determine if plant life strategies mediate differences in plant communities in response to great skuas, data were reanalysed using a Correlative Response Model (CRM) implemented in the BORAL package in R (*53*). Plant strategies (competitor (C), stress-tolerant (S) and ruderal (R)) were assigned according to Grime et al. (*31*), and the associated scores on a scale of 0-4 representing the strength of CSR characteristics of all species used as a response variable (*54*). The CRM fits separate GLMs for each species abundance using a common set of explanatory variables by Bayesian MCMC estimation. To account for residual covariation between species abundances not explained by the specified explanatory variables, such as biotic interactions, two unmeasured latent variables were included in the model. Species-specific traits were incorporated into the model as random effects and the degree to which a trait is associated with the explanatory variable is reflected in the model coefficients (*53,55*). In all models the number of pin touches for each plant species was used as a response variable and great skua density (individual sampling location counts) and C-S-R scores were incorporated as random effects. The association of a plant life strategy with great skua density was considered significant if the 95% credible intervals of the trait coefficient did not include zero (*53*). All models were verified using Dunn-Smyth residual analysis (*53*).

The effect of great skua density on carabid species richness was assessed using mixed effects models fit using lme4. Great skua density was included as a fixed effect and colony (Fetlar, Hermaness, Noss) as a random effect. The effect of great skuas on terrestrial invertebrate order and carabid species abundance was assessed using generalized linear models (GLM) implemented through the MVAbund package, in R (*52*). Here, the response variables comprised either the number of individuals in each invertebrate order or the number of individuals in each carabid species at each sampling location (i.e. a site by order/species matrix). In both models, great skua density was used as explanatory variables with ‘site’ as a random factor. Score tests (a resampling-based hypothesis test) using PIT-trap resampling were then used to initially make community-level (i.e. all taxa abundances at a sample location combined) and, subsequently, taxon-specific inferences about how the explanatory variables influenced the invertebrate order and carabid species abundances. This test and the resampling technique were considered most appropriate for these data structures, whilst still taking into account correlations between taxa abundance (*52*).

A structural equation model (SEM) was fitted to standardized data (zeta scores or, in other words, the correlation matrix; *56-58*). All the various forms of soil nutrients were considered in the construction of the latent “soil nutrients” but those presented returned the best fitting and most parsimonious models. The presented path structure is based on the general hypothesis that great skua density impacts the biological communities (plants and carabids) both directly and indirectly via soil nutrients, and both direct and indirect paths were supported by the fitted model and thus retained in the finally presented model. We found little support for model including direct effect of location variables (latitude, longitude and functions of latitude and longitude) and these were therefore excluded from the final models (in other words, SEM with location variables returned poor global fit metrics and were rejected at a Chi-square test set at the conservative *P* value of 0.05). We found no literature evidence and had no specific hypothesis on causal directionality of the correlation observed between carabid richness and soil nutrients, and we therefore modelled that correlation as a non-directional covariance term.

#### Transplant experiments

The change in plant community composition in response to increasing and decreasing great skua density was analysed and visualised using principal coordinates analysis (PCoA) as in the stratified surveys. Plant nutrient status (nitrogen and phosphorus) response to increasing and decreasing great skua densities was assessed using linear models with treatment as the only predictor variable. Where turfs were exposed to increased great skua density treatments comprised high to high skua density control, low to high skua density treatment and low to low skua density control; and where turfs were exposed to decreasing great skua density treatments comprised high to high skua density control, high to low skua density treatment and low to low skua density control. A treatment effect was initially assessed using ANOVA and differences between the treatments was assessed using post-hoc Least Significant Difference (LSD) tests.

## Acknowledgments

We thank NatureScot, the Royal Society for the Protection of Birds, Richard D Bardgett, Ross Barnett, Juan Brown, Andrew Denton, Martha Devine, Chris Dosser, Ully Kritzler, Ben Leonard, Craig Nisbet, and Jonathan Swale.

## Funding

Macaulay Development Trust University of Stirling studentship Natural Environment Research Council

Natural Environment Research Council National Environmental Isotope Facility

## Author contributions

Conceptualization: LG, DJ

Funding acquisition: LG, DJ, KP

Methodology: LG, DJ

Investigation: MG

Formal Analysis: MG, TC

Supervision: LG, DJ, KP

Writing – original draft: DJ, LG

Writing – editing and reviewing: DJ, LG, MG, TC, KP

## Data and materials availability

Contact corresponding authors

## Supporting Information

**Fig. S1.**
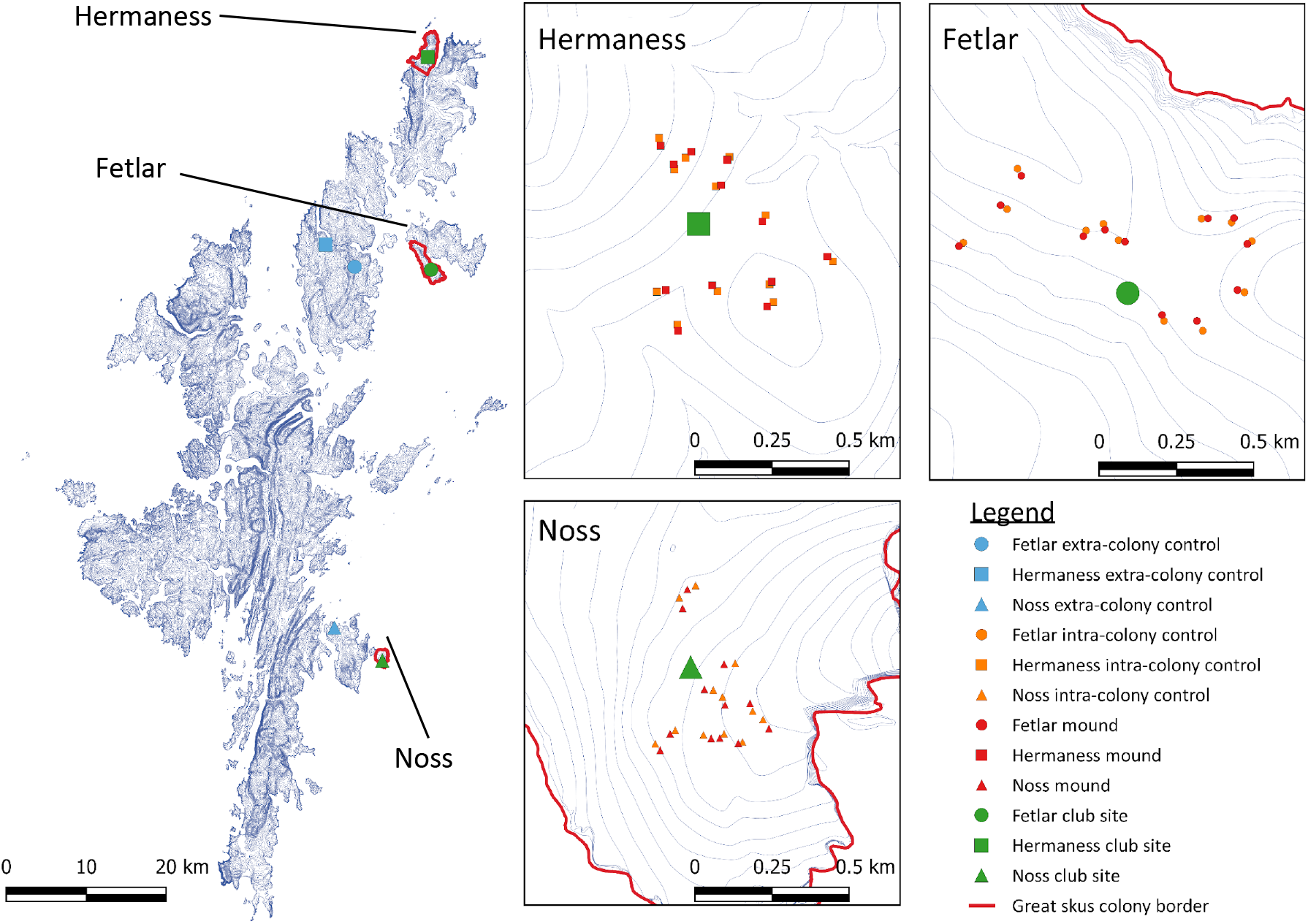
The correlative study design. In the map of Shetland (left) the large green points represent the high great skua density club sites and the large blue points the low great skua density sites outside the colonies. In the individual colony maps, the small red and orange points represent high great skua density mounds and low great skua density colony locations, and the large green points represent the high great skua density club site locations. The shape represents the three separate colonies (circle: Fetlar, square: Hermaness, and triangle: Noss).

**Fig. S2.**
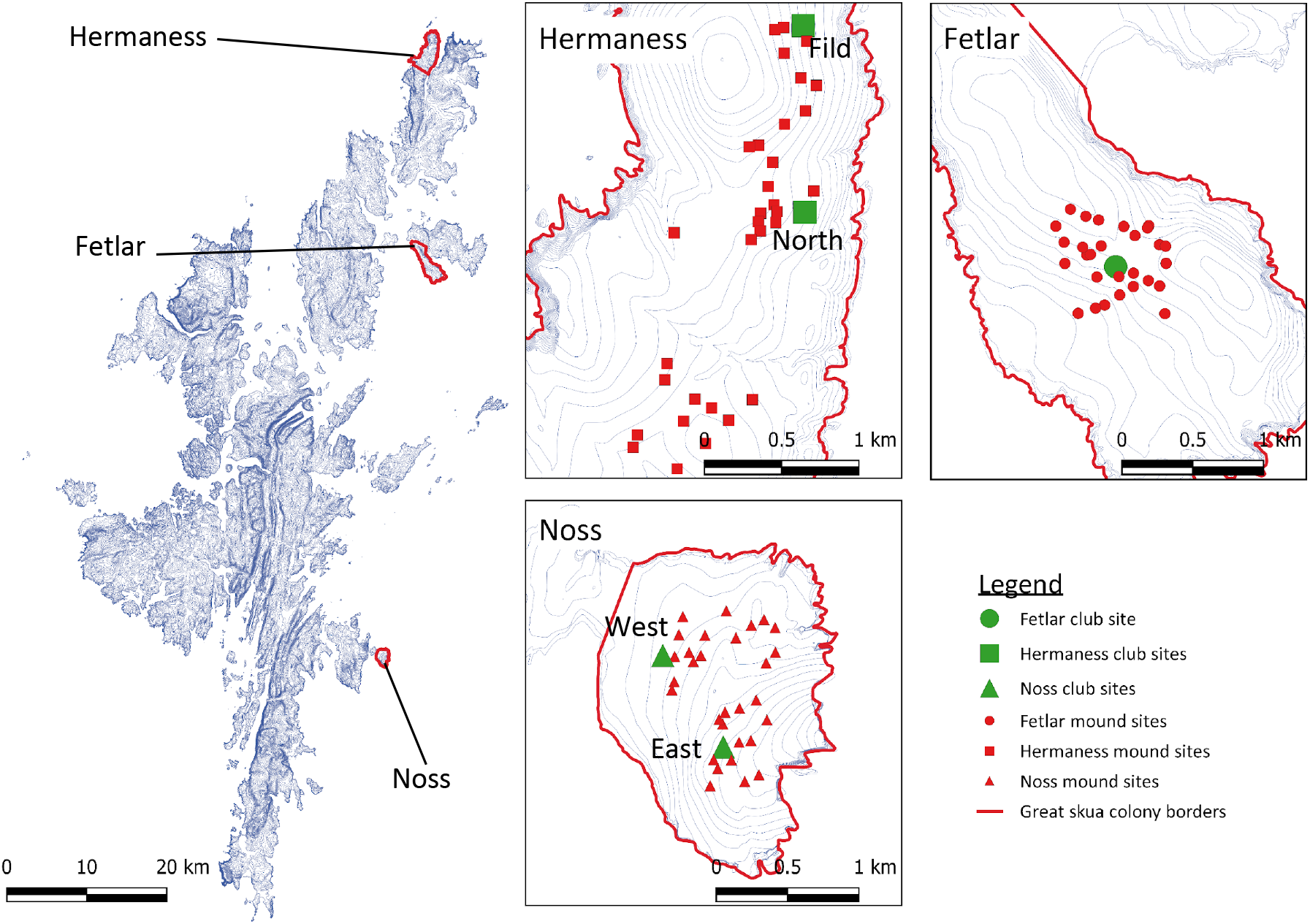
A map of the study design used to determine the amount of nutrients deposited by great skua into the terrestrial ecosystem. The large green symbols identify the location of each high great skua density club site, with the name of that club sites appear in the text adjacent, where there is more than one. The small red symbols identify the location of the high great skua density mound sampling sites. Low great skua density sites not shown.

**Fig. S3.**
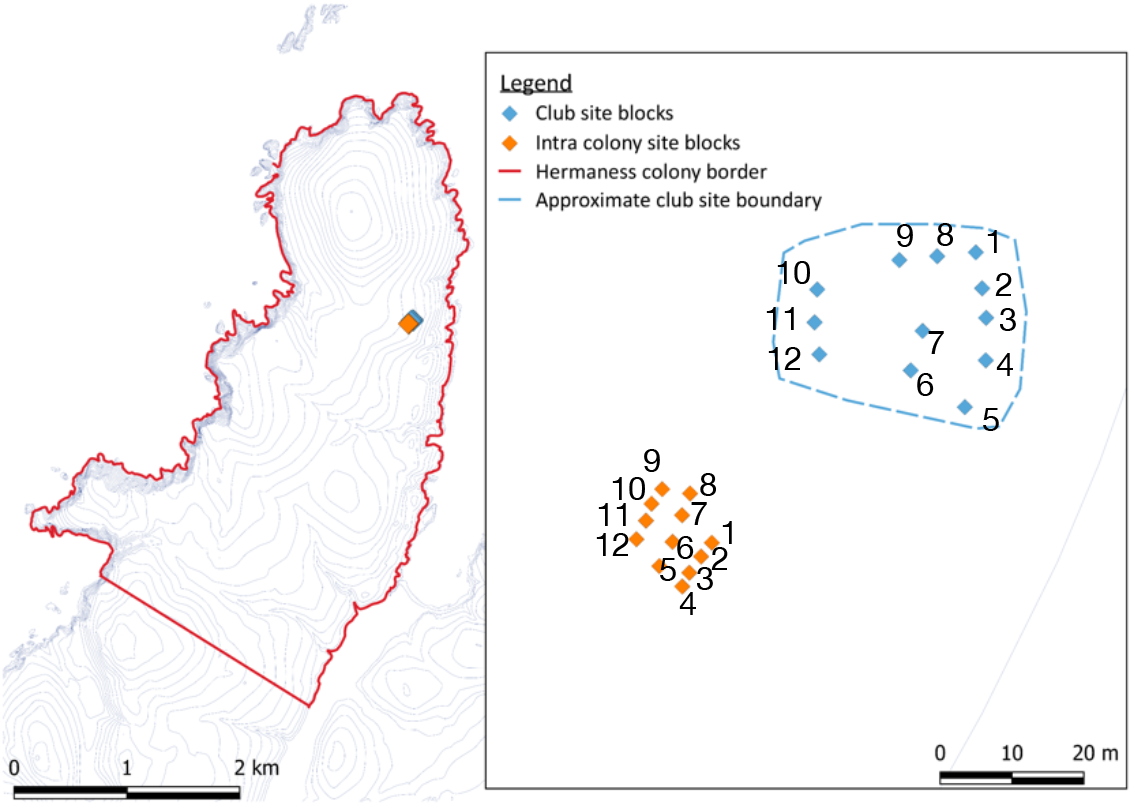
The reciprocal turf transplant study design and location at the Hermaness great skua colony. In the map on the right, the blue points represent the blocks in the high great skua density club site and the orange points represent blocks in the low great skua density colony site. At each block, two turfs were dug, one of which was transplanted to the paired block (same numbers) in the opposing site and the other was placed back into the site of origin.

**Fig. S4.**
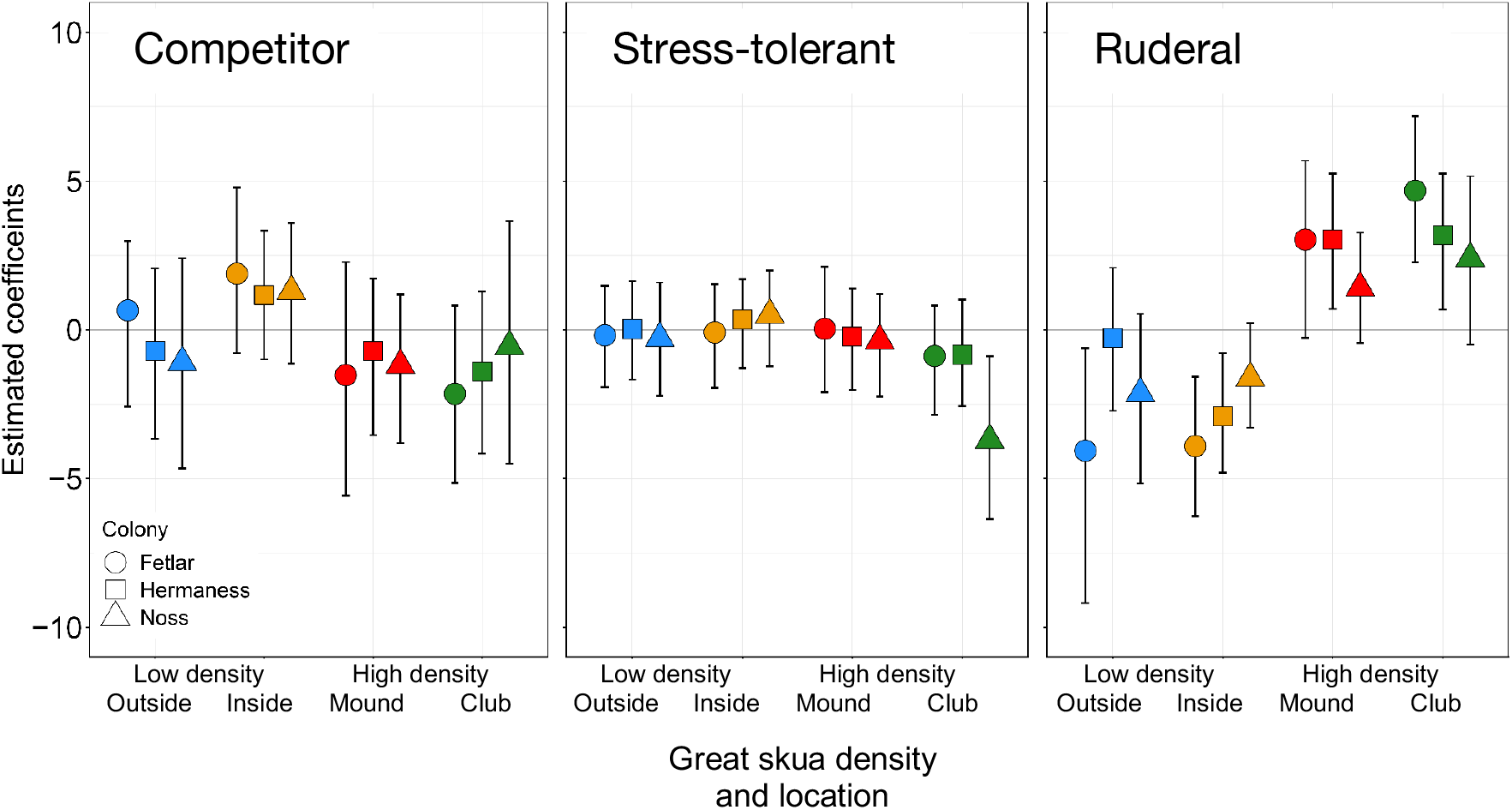
Association between great skua density (low density outside colony; low density inside colony; high density mound; high density club site) and life strategy (competitor, C; stress tolerant, S; ruderal, R) of associated plants in each colony. Each point represents an estimated coefficient of each of the each of the C-S-R scores (derived from Grime et al., 2007) as estimated by the correlative response model. The error bars represent 95% credible intervals. Where the error bars do not overlap with zero there is a significant association between the life strategy and great skua density.

**Fig. S5.**
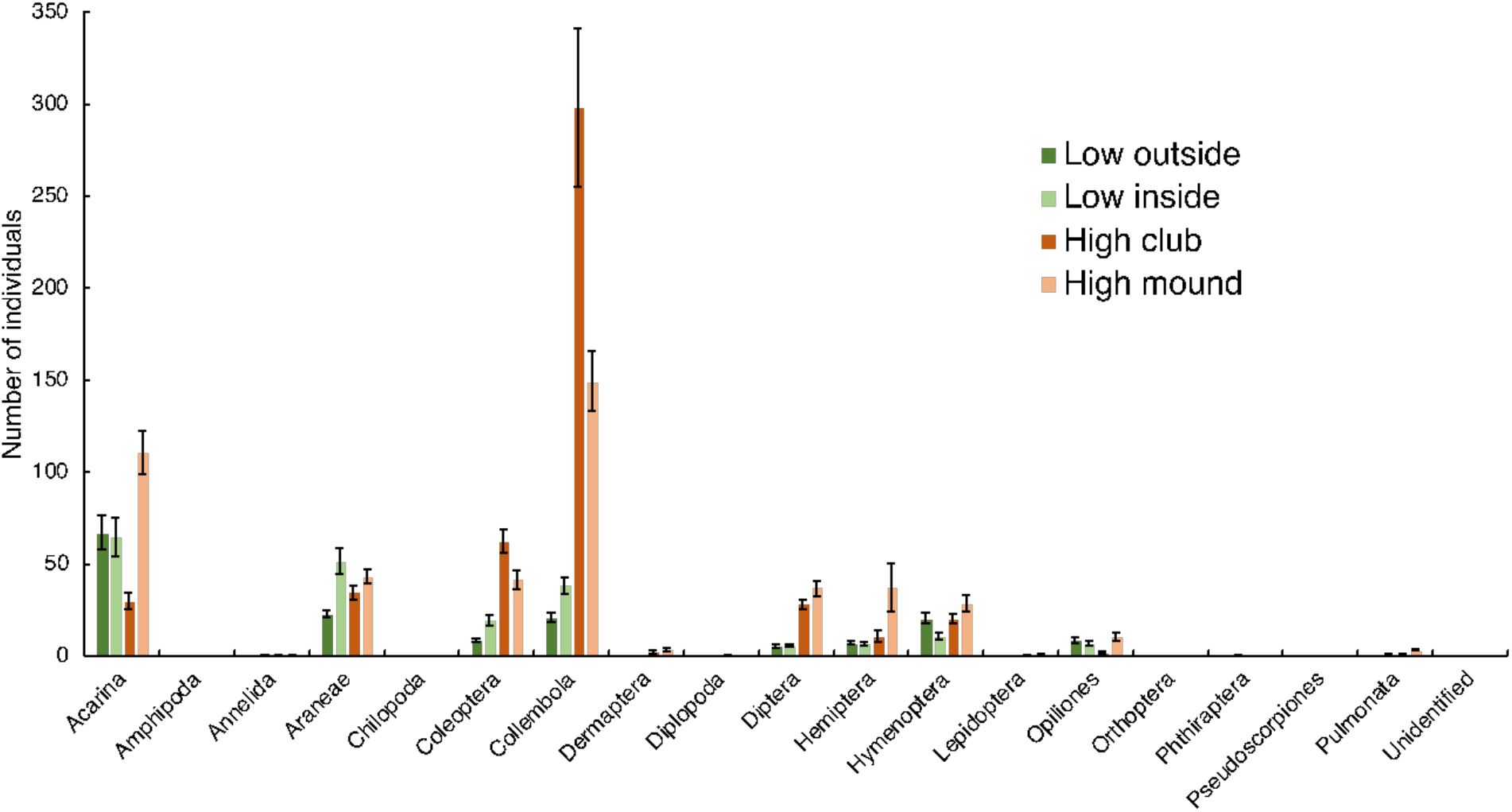
Number of individuals of invertebrates recovered from pitfall traps grouped by taxonomic order in relation to great skua density site category (high density club and mound; low density inside and outside colony) (means ± SEM). Refer to Table S2 for statistical outputs.

**Fig. S6.**
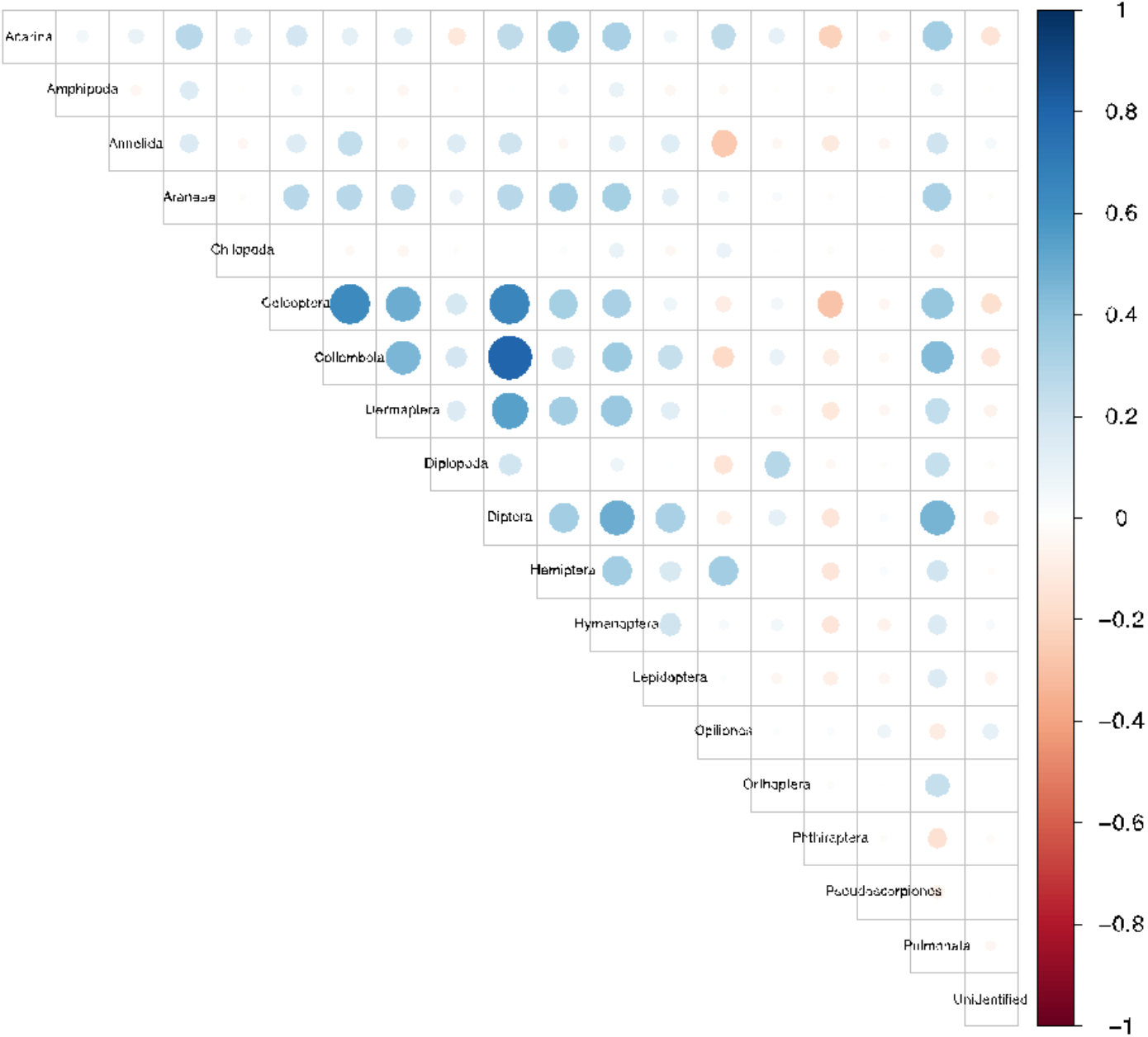
Correlation matrix between the abundance of invertebrates within taxonomic orders. Positive associations are shown in blue and negative associations in red.

**Fig. S7.**
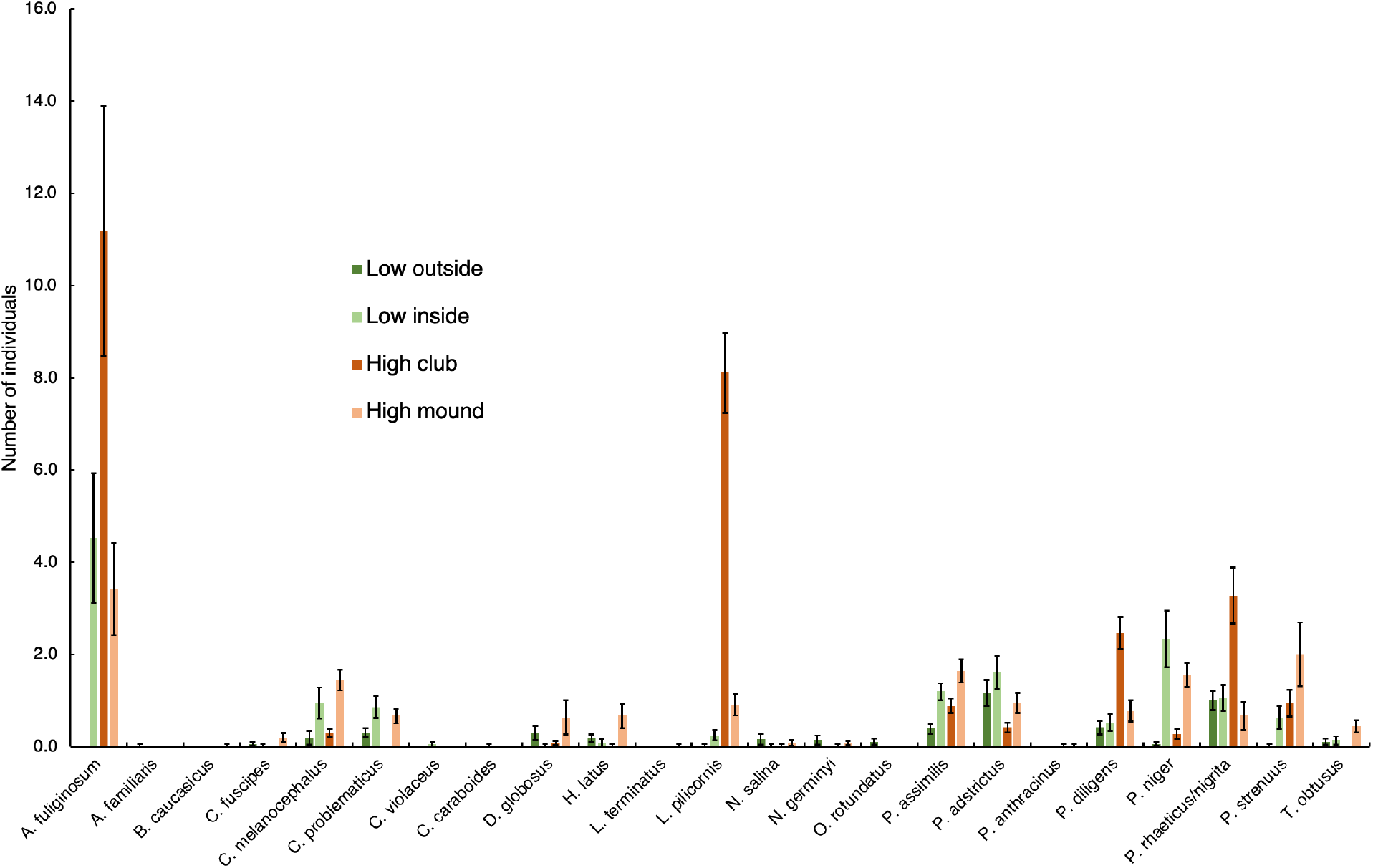
Number of individuals of carabids (ground beetles) recovered from pitfall traps grouped by species in relation to great skua density site category (high density club and mound; low density inside and outside colony) (means ± SEM). Genus abbreviation letters refer to: *Agonum, Bradycellus, Calathus, Dyschirius, Harpalus, Loricera, Nebria, Olisthopus, Pterostichus, Trechus.* Refer to Table S3 for statistical outputs and full genus names.

**Table S1.** The effect of great skua density site category (high density club and mound; low density inside and outside colony), colony and the interaction on plant species abundance in the correlative study. C, S and R refers to the plant life strategy (Competitor, Stress-tolerant and Ruderal) classification triangle (Grime 1979, 2007) as per fig. S4. The row colours reflect groups of species that associate with exclusively low great skua density areas (blue), exclusively high great skua density club sites (green), and both high (mounds) and low (inside colony) great skua density areas (red) (these were on Noss). Significant effects are denoted by bold text and non-significant effects by ns.

| Species | Strategy | Great skua density |  | Colony |  | Interaction |  |
| --- | --- | --- | --- | --- | --- | --- | --- |
|  |  | Score | P | Score | P | Score | P |
| *Agrostis capillaris* | CSR | 3.808 | ns | 2.826 | ns | 0.001 | ns |
| *Aira praecox* | RS | 3.824 | ns | 1.426 | ns | 0.338 | ns |
| *Carex bigelowii* | No data | 2.083 | ns | 1.482 | ns | 0.001 | ns |
| *Carex binervis* | S | 1.579 | ns | 2.455 | ns | 1.773 | ns |
| *Carex panicea* | S/SC | 2.370 | ns | 2.635 | ns | 2.109 | ns |
| *Carex pilulifera* | S | 3.000 | ns | 2.000 | ns | 0.001 | ns |
| *Dactylorhiza maculata* | SR | 2.180 | ns | 1.121 | ns | 2.326 | ns |
| *Drosera rotundifolia* | SR | 1.352 | ns | 0.925 | ns | 0.001 | ns |
| *Erica cinerea* | S/SC | 5.634 | ns | 4.993 | ns | 3.897 | ns |
| *Eriophorum vaginatum* | S/SC | 3.674 | ns | 4.555 | ns | 12.68 | ns |
| *Euphrasia officinalis agg.* | SR | 2.490 | ns | 3.152 | ns | 0.001 | ns |
| *Juncus effusus* | C/SC | 1.724 | ns | 1.182 | ns | 0.001 | ns |
| *Juncus squarrosus* | S/CSR | 2.378 | ns | 0.203 | ns | 4.821 | ns |
| *Narthecium ossifragum* | S/CSR | 2.565 | ns | 3.866 | ns | 2.728 | ns |
| *Pinguicula vulgaris* | SR | 3.480 | ns | 2.459 | ns | 0.001 | ns |
| *Polygala serpyllifolia* | S/CSR | 2.234 | ns | 2.855 | ns | 0.001 | ns |
| *Rumex acetosa* | CSR | 1.741 | ns | 1.732 | ns | 0.133 | ns |
| *Rumex acetosella* | SR/CSR | 5.968 | ns | 4.318 | ns | 0.001 | ns |
| *Sagina nodosa* | S/SR | 0.608 | ns | 0.413 | ns | 0.001 | ns |
| *Succisa pratensis* | S/CSR | 1.698 | ns | 1.183 | ns | 0.001 | ns |
| *Vaccinium myrtillus* | SC | 3.970 | ns | 1.101 | ns | 2.767 | ns |
| *Viola sp.* | No data | 1.686 | ns | 4.513 | ns | 0.000 | ns |
| *Calluna vulgaris* | S/SC | 24.39 | **0.001** | 19.88 | **0.002** | 36.90 | **0.015** |
| *Empetrum nigrum* | CS | 8.358 | **0.019** | 20.58 | **0.001** | 19.44 | ns |
| *Erica tetralix* | S/SC | 14.01 | **0.003** | 4.046 | ns | 8.235 | ns |
| *Eriophorum angustifolium* | S/SC | 32.20 | **0.001** | 7.613 | ns | 27.93 | **0.024** |
| *Trichophorum cespitosum* | S/SC | 19.55 | **0.001** | 0.512 | ns | 8.739 | ns |
| *Neottia cordata* | No data | 9.267 | **0.019** | 5.463 | ns | 0.000 | ns |
| *Anthoxanthum odoratum* | SR/CSR | 27.88 | **0.001** | 10.10 | **0.038** | 29.43 | **0.024** |
| *Carex nigra* | S/SC | 14.93 | **0.003** | 15.40 | **0.002** | 19.12 | ns |
| *Cerastium fontanum* | R/CSR | 20.67 | **0.001** | 0.966 | ns | 2.761 | ns |
| *Deschampsia flexuosa* | S/SC | 14.54 | **0.003** | 12.65 | **0.006** | 11.98 | ns |
| *Galium saxatile* | S/CSR | 19.21 | **0.001** | 8.154 | ns | 10.43 | ns |
| *Luzula multiflora* | S/CSR | 38.54 | **0.001** | 9.926 | **0.038** | 17.47 | ns |
| *Molinia caerulea* | SC | 8.152 | **0.019** | 5.128 | ns | 5.644 | ns |
| *Nardus stricta* | S/SC | 7.390 | **0.02** | 4.398 | ns | 120.7 | **0.015** |
| *Potentilla erecta* | S/CSR | 14.29 | **0.003** | 30.61 | **0.001** | 16.26 | ns |
| *Agrostis stolonifera* | CR | 16.60 | **0.002** | 7.683 | ns | 15.16 | ns |
| *Holcus lanatus* | CSR | 7.443 | **0.02** | 17.03 | **0.002** | 9.462 | ns |
| *Juncus bufonius* | R/SR | 11.47 | **0.008** | 2.492 | ns | 0.001 | ns |
| *Poa pratensis* | CSR | 8.691 | **0.019** | 6.830 | ns | 0.001 | ns |
| *Sagina procumbens* | R/CSR | 7.910 | **0.019** | 1.841 | ns | 2.896 | ns |
| *Stellaria media* | R | 23.63 | **0.001** | 7.107 | ns | 4.888 | ns |

**Table S2.** The effects of great skua density site category (high density club and mound; low density inside and outside colony), colony and their interaction on the abundance of invertebrates within taxonomic orders. The significance of the predictor is calculated using ANOVA on 999 bootstrap resampling iterations. Significant effects are denoted by bold text, and non-significant effects by ns.

| Invertebrate order | Great skua density |  | Colony |  | Interaction |  |
| --- | --- | --- | --- | --- | --- | --- |
|  | Score | *P* | Score | *P* | Score | *P* |
| Acarina | 35.81 | **0.001** | 40.18 | **0.001** | 35.71 | **0.011** |
| Amphipoda | 3.000 | ns | 2.000 | ns | 0.001 | ns |
| Annelida | 5.116 | ns | 14.32 | **0.025** | 4.903 | ns |
| Araneae | 21.33 | **0.001** | 16.30 | **0.012** | 49.54 | **0.004** |
| Chilopoda | 3.000 | ns | 2.000 | ns | 0.001 | ns |
| Coleoptera | 58.97 | **0.001** | 36.74 | **0.001** | 45.93 | **0.004** |
| Collembola | 80.73 | **0.001** | 14.98 | **0.02** | 34.31 | **0.012** |
| Dermaptera | 20.14 | **0.001** | 12.52 | **0.029** | 5.657 | ns |
| Diplopoda | 5.001 | ns | 6.228 | ns | 0.001 | ns |
| Diptera | 62.71 | **0.001** | 36.53 | **0.001** | 39.24 | **0.005** |
| Hemiptera | 61.32 | **0.001** | 8.763 | ns | 28.06 | **0.022** |
| Hymenoptera | 17.97 | **0.001** | 33.51 | **0.001** | 23.28 | **0.03** |
| Lepidoptera | 19.45 | **0.001** | 12.37 | **0.029** | 3.769 | ns |
| Opiliones | 16.13 | **0.003** | 17.32 | **0.009** | 49.68 | **0.004** |
| Orthoptera | 1.724 | ns | 1.182 | ns | 0.001 | ns |
| Phthiraptera | 4.197 | ns | 9.528 | ns | 0.000 | ns |
| Pseudoscorpiones | 3.000 | ns | 2.000 | ns | 0.001 | ns |
| Pulmonata | 42.41 | **0.001** | 11.30 | **0.044** | 5.709 | ns |
| Unidentified | 6.000 | ns | 4.000 | ns | 0.001 | ns |

**Table S3.** The effect of great skua density site category (high density club and mound; low density inside and outside colony), colony and their interaction on carabid (ground beetle) species abundance. The significance of the predictor is calculated using ANOVA on 999 bootstrap resampling iterations. Significant effects are denoted by bold text and non-significant effects by ns.

| Carabid species | Great skua density |  | Colony |  | Interaction |  |
| --- | --- | --- | --- | --- | --- | --- |
|  | Score | *P* | Score | *P* | Score | *P* |
| *Agonum fuliginosum* | 495.9 | **0.001** | 1233 | **0.001** | 7.62 | ns |
| *Amara familiaris* | 3.00 | ns | 2.00 | ns | 0.00 | ns |
| *Bradycellus caucasicus* | 3.00 | ns | 2.00 | ns | 0.00 | ns |
| *Calathus fuscipes* | 10.46 | ns | 7.82 | ns | 6.81 | ns |
| *Calathus melanocephalus* | 39.52 | **0.014** | 13.03 | ns | 28.53 | ns |
| *Carabus problematicus* | 28.60 | **0.025** | 1.81 | ns | 14.51 | ns |
| *Carabus violaceus* | 6.00 | ns | 4.00 | ns | 0.00 | ns |
| *Cychrus caraboides* | 3.00 | ns | 2.00 | ns | 0.00 | ns |
| *Dyschirius globosus* | 30.71 | **0.025** | 29.46 | **0.018** | 32.72 | ns |
| *Harpalus latus* | 37.51 | **0.017** | 53.20 | **0.001** | 35.91 | ns |
| *Leistus terminatus* | 3.00 | ns | 2.00 | ns | 0.00 | ns |
| *Loricera pilicornis* | 697.1 | **0.001** | 8.69 | ns | 32.96 | ns |
| *Nebria salina* | 6.09 | ns | 6.73 | ns | 11.00 | ns |
| *Notiophilus biguttatus* | 3.00 | ns | 2.00 | ns | 0.00 | ns |
| *Notiophilus germinyi* | 15.00 | ns | 9.39 | ns | 13.00 | ns |
| *Olisthopus rotundatus* | 12.00 | ns | 3.50 | ns | 0.00 | ns |
| *Patrobus assimilis* | 27.62 | **0.025** | 4.04 | ns | 25.63 | ns |
| *Pterostichus adstrictus* | 25.67 | **0.025** | 26.36 | **0.018** | 36.92 | ns |
| *Pterostichus anthracinus* | 2.00 | ns | 1.00 | ns | 2.00 | ns |
| *Pterostichus diligens* | 96.99 | **0.001** | 7.84 | ns | 52.58 | **0.043** |
| *Pterostichus niger* | 119.0 | **0.001** | 87.41 | **0.001** | 9.81 | ns |
| *Pterostichus rhaeticus/nigrita* | 102.1 | **0.001** | 190.6 | **0.001** | 10.12 | ns |
| *Pterostichus strenuus* | 81.39 | **0.001** | 242.3 | **0.001** | 68.81 | **0.043** |
| *Trechus obtusus* | 18.59 | **0.043** | 12.48 | ns | 8.76 | ns |

**Table S4.** Mean (± SEM) nutrient concentrations (%) and ratios in plant litter collected from high great skua density club sites and low great skua density sites outside skua colonies used in the decomposition experiment. Values sharing a letter within columns are not significantly different (*P* >0.05). Nutrients are carbon (C), nitrogen (N) and phosphorus (P).

| Great skua density | C (%) | N (%) | P (mg g<sup>-1</sup> dwt) | C:N | C:P | $\delta^{15}\text{N}$ ‰ |
| --- | --- | --- | --- | --- | --- | --- |
| High | 45.3<sup>a</sup> ± 0.19 | 1.39<sup>a</sup> ± 0.013 | 1.19<sup>a</sup> ± 0.069 | 32.6<sup>a</sup> ± 0.30 | 395<sup>a</sup> ± 25.8 | 3.30<sup>a</sup> ± 0.112 |
| Low | 45.2<sup>a</sup> ± 0.33 | 0.89<sup>b</sup> ± 0.012 | 0.45<sup>b</sup> ± 0.005 | 50.8<sup>b</sup> ± 0.48 | 1011<sup>b</sup> ± 12.0 | -0.49<sup>b</sup> ± 0.107 |

**Table S5.** Mean phosphorus concentrations (mg g^-1^ dwt ± SEM) in leaves of plants in the turf transplant experiment. Low to low and high to high are control turfs that were extracted and replaced in their original locations. Values sharing a letter within a treatment group and species are not significantly different (*P* >0.05).

| Species | Increasing great skua density |  |  | Decreasing great skua density |  |  |
| --- | --- | --- | --- | --- | --- | --- |
|  | Low to low | Low to high | High to high | High to high | High to low | Low to low |
| *P. erecta* | 3.78<sup>a</sup> ± 0.43 | 4.04<sup>a</sup> ± 0.74 | 5.15<sup>a</sup> ± 0.37 | 5.15<sup>b</sup> ± 0.37 | 4.60<sup>ab</sup> ± 0.21 | 3.78<sup>a</sup> ± 0.37 |
| *A. odoratum* | 2.69<sup>a</sup> ± 0.44 | 3.10<sup>a</sup> ± 0.26 | 3.59<sup>a</sup> ± 0.38 | 3.59<sup>b</sup> ± 0.38 | 5.18<sup>a</sup> ± 0.32 | 2.69<sup>b</sup> ± 0.44 |
| *E. vaginatum* | 1.50<sup>a</sup> ± 0.45 | 1.79<sup>ab</sup> ± 0.32 | 2.68<sup>b</sup> ± 0.22 | 2.68<sup>a</sup> ± 0.22 | 1.81<sup>ab</sup> ± 0.27 | 1.50<sup>a</sup> ± 0.45 |
| *C. vulgaris* | 1.15<sup>a</sup> ± 0.18 | 1.95<sup>b</sup> ± 0.16 | 2.11<sup>b</sup> ± 0.13 | 2.11<sup>b</sup> ± 0.13 | 2.38<sup>b</sup> ± 0.17 | 1.15<sup>a</sup> ± 0.18 |

**Table S6.** Full output from the structural equation model (see Fig. 1J). SEM = standard error of mean; In the regressions, the dependent variable is in *italics*.

| | Estimate | SEM | z-value | $P(>\vert z\vert )$ | Std. lv | Std. all |
| --- | --- | --- | --- | --- | --- | --- |
| **Latent variables** |  |  |  |  |  |  |
| NH<sub>4</sub> | 0.390 | 0.074 | 5.267 | 0.000 | 0.465 | 0.467 |
| PO<sub>4</sub> | 0.703 | 0.076 | 9.217 | 0.000 | 0.839 | 0.843 |
| N | 0.495 | 0.072 | 6.867 | 0.000 | 0.591 | 0.593 |
| **Regressions** |  |  |  |  |  |  |
| *Soil nutrients~* |  |  |  |  |  |  |
| Great skua density | 0.653 | 0.117 | 5.593 | 0.000 | 0.547 | 0.545 |
| *Plant community axis~* |  |  |  |  |  |  |
| Great skua density | 0.181 | 0.088 | 2.052 | 0.040 | 0.181 | 0.181 |
| Soil nutrients | 0.495 | 0.080 | 6.155 | 0.000 | 0.590 | 0.593 |
| *Plant species richness~* |  |  |  |  |  |  |
| Plant community axis | 0.858 | 0.109 | 7.899 | 0.000 | 0.858 | 0.857 |
| Great skua density | -0.233 | 0.086 | -2.716 | 0.007 | -0.233 | -0.233 |
| Soil nutrients | -0.279 | 0.108 | -2.581 | 0.010 | -0.333 | -0.335 |
| *Carabid species richness~* |  |  |  |  |  |  |
| Plant community axis | 0.355 | 0.098 | 3.636 | 0.000 | 0.355 | 0.355 |
| Plant species richness | 0.200 | 0.079 | 2.525 | 0.012 | 0.200 | 0.200 |
| **Covariances** |  |  |  |  |  |  |
| *Soil nutrients~~* |  |  |  |  |  |  |
| Carabid species richness | 0.234 | 0.097 | 2.401 | 0.016 | 0.234 | 0.288 |
| **Variances** |  |  |  |  |  |  |
| NH<sub>4</sub> | 0.776 | 0.098 | 7.910 | 0.000 | 0.776 | 0.782 |
| PO<sub>4</sub> | 0.287 | 0.081 | 3.559 | 0.000 | 0.287 | 0.290 |
| N | 0.643 | 0.087 | 7.381 | 0.000 | 0.643 | 0.648 |
| Plant community axis | 0.494 | 0.074 | 6.650 | 0.000 | 0.494 | 0.498 |
| Plant species richness | 0.608 | 0.076 | 7.952 | 0.000 | 0.608 | 0.613 |
| Carabid species richness | 0.661 | 0.081 | 8.156 | 0.000 | 0.661 | 0.666 |
| **Variable** | **R<sup>2</sup>** |  |  |  |  |  |
| NH<sub>4</sub> | 0.218 |  |  |  |  |  |
| PO<sub>4</sub> | 0.710 |  |  |  |  |  |
| N | 0.352 |  |  |  |  |  |
| Plant community axis | 0.502 |  |  |  |  |  |
| Plant species richness | 0.387 |  |  |  |  |  |
| Carabid species richness | 0.334 |  |  |  |  |  |
| Soil nutrients | 0.298 |  |  |  |  |  |

